# The Rising Prevalence of Weapons in Unsafe Arming Configurations Discovered in American Airports: the Increasing Practice of Storage and Carry of Firearms with a Round Chambered

**DOI:** 10.1101/613687

**Authors:** Sherry Towers, Bechir Amoundi, Richard Cordova, Karen Funderburk, Cesar Montalvo, Mugdha Thakur, Josean Velazquez-Molina, Carlos Castillo-Chavez

## Abstract

**Background:** Recent studies have shown that Americans appear to be increasingly owning and carrying firearms for personal protection, and are increasingly storing their firearms loaded. However, the prevalence of firearm carry and/or storage with a round chambered has not hitherto been studied; in this arming configuration the weapon is loaded, with a bullet resting against the firing pin, ready to fire if the trigger is pulled. Ostensibly this decreases reaction time in a threat situation, but carries an increased risk of accidental discharge. It is unknown what fraction of firearm owners typically carry or store their firearms in this configuration, and if there have been significant temporal trends in the practice.

**Objective:** We examine firearms detected at Transportation Security Administration (TSA) airport security checkpoints between 2012 to 2017, to examine geospatial and temporal trends in the prevalence of unsafe arming configuration in detected firearms.

**Results:** The fraction of detected firearms found to be loaded has risen significantly since 2012 (Beta Binomial logistic regression *p* = 0.011). States with firearm child access prevention laws have significantly fewer firearms found by the TSA to be loaded (*p* = 0.039).

The fraction of loaded firearms found by the TSA to also have a round chambered has also risen significantly since 2012 (Beta Binomial logistic regression *p* < 0.001). By 2017, 36% of firearms found loaded were also found to have a round chambered, representing an apparent concerning trend in unsafe arming configurations during carry and storage.

**Conclusions:** Americans appear to be increasingly using and storing firearms in unsafe arming configurations. TSA firearm detection data can potentially provide a key source of data when researching trends in firearm injury.

## 1 Introduction

Firearms are the third leading cause of injury related deaths in the United States [1], and America has higher rates of intentional and accidental firearm injury than any other developed country [2, 3]. Over 90% of firearm deaths of children 0 to 14 worldwide occur in the United States [3, 4]; each year nearly 1,300 children under the age of 18 die and 5,790 are treated for gunshot wounds [4]. Firearm related deaths are the third leading cause of death overall among U.S. children aged 1 to 17 years [4], and the fourth leading cause of death among children aged 1-14 years [5].

While high prevalence of firearm ownership in the US has been shown to play a role in these patterns [6–10], unsafe storage practices of firearms, such as storing firearms loaded, have been associated with additional increased risk of firearm injury and mortality [10–14], particularly among pediatric populations [15–20]. Beyond the issue of storage however, there has been a dearth of studies on the prevalence of various unsafe carrying practices, and their potential impact on accidental shootings. Carrying practices that might increase the risk of such incidents include carrying with the safety off, without a holster, or with a round chambered. In this arming configuration, a round of ammunition is chambered in the firearm, resting against the firing pin, and the firearm is at most two steps from being fired; turning off the safety if there is one (and it is on), and pulling the trigger. In general, firearm safety training discourages the practice of storing or carrying a firearm with a round chambered because of the increased risk of accidental injury. To the authors’ knowledge, no study has yet estimated the prevalence of this practice among American firearm owners, despite the additional risks it likely poses to public health.

Understanding the role that unsafe storage and carrying practices play in firearm injury and mortality requires temporal and geospatial estimates of the prevalence of these practices. Prior research on firearm storage habits has often been based on national survey data collected by the annual Behavioral Risk Factor Surveillance System (BRFSS), conducted by the U.S. Centers for Disease Control (CDC) (for example, References [8, 13, 21]). However, questions on this survey surrounding firearm habits ceased in 2004. Other national surveys of storage and carrying practices are expensive to undertake and are thus often limited in size, such that the results do not necessarily provide reliable estimates at the state or local level, and may also suffer from recall and/or social desirability bias [22, 23].

Here, we examine firearms detected at airport security checkpoints as a potential means to estimate the temporal and geospatial trends in the prevalence of unsafe storage and carry practices. Since October, 2011, the Transportation Security Administration (TSA) has made public on its weekly blog tallies of the number of firearms detected at airport passenger security checkpoints (www.tsa.gov/blog, accessed March, 2019). Since January, 2012, the recorded information contains the date the firearm was found, the airport at which it was found, the caliber of the firearm, and whether or not it was loaded. The TSA data also indicate whether or not a round of ammunition was chambered in the firearm.

The TSA states that most passengers found with a firearm claim that they had simply forgotten that the firearm was stored in their luggage, and had not noticed it when they packed (see, for example, https://bit.ly/2Bu1JG7, accessed March, 2019). However, it is quite possible that some of these passengers were in truth intentionally carrying a firearm due to ignorance of federal laws (or even potential malintent), and had simply claimed they had forgotten the firearm to try to avoid legal penalties.

The fraction of passengers found to have firearms at airport security checkpoints is thus influenced by several factors:

1. The prevalence of firearm ownership.
2. The prevalence of firearm owners storing firearms in luggage.
3. The fraction of firearms stored in luggage not noticed by people packing.
4. The prevalence of firearm carry.
5. The prevalence, among those who carry, of ignorance of federal laws forbidding firearms on airplanes.
6. The efficiency of the TSA in detecting firearms.

The first, second, and fourth factors may have both geospatial and temporal trends, yet the third and fifth are likely to be universal across time and locale. The last factor is difficult to estimate; in 2015, a Department of Homeland Security (DHS) study showed that TSA screening only had 5% efficiency in detecting prohibited items at airport security checkpoints, but since that time screening methods have likely improved (see https://bit.ly/2ocyd1O, accessed March, 2019). However, TSA screening methods are standardised across the U.S., thus the fraction of the total number of passengers from 2012 onwards found to have firearms is reflective, at the state level and down to a scale factor, of geospatial differences in the prevalence of a combination of unsafely stored firearms and firearm carry.

It is important to note that the fraction of detected firearms found loaded is independent of the TSA firearm detection efficiency, since that efficiency equally affects both the numerator and denominator. Similarly, the fraction of loaded firearms also found to have a round chambered is also independent of TSA firearm detection efficiencies. Both of these quantities can thus be used to estimate trends in unsafe arming configuration practices during storage and carry among a cross-section of firearm owners.

In the following sections we describe our sources of data and statistical methodologies used in the temporal and geospatial analyses, followed by a presentation of results and discussion.

## 2 Methods and Materials

### 2.1 Data

#### 2.1.1 TSA firearm detection data

Since October, 2011 the TSA has maintained a website providing weekly updates on the prohibited items discovered at TSA airport passenger security passenger checkpoints (available at www.tsa.gov/blog, accessed March, 2019). The data include the details of individual incidents of firearms detected, including the airport at which each firearm was detected, the date it was found, the caliber of the firearm, whether it was loaded, and whether it also had a round chambered. It was only after partway through December, 2011, however, that the TSA listed the details of unloaded firearms in addition to the loaded firearms. From this data, we compiled the list of 14, 866 firearms detected between 2012 to 2017. A total of 12, 711 (85.5%) were found loaded, and 4, 493 (30.2%) also had a round chambered.

#### 2.1.2 Airline passenger data

To estimate the number of passengers screened by the TSA at airports across the U.S. in order to estimate per-passenger geospatial trends in TSA firearm detections, we compiled airport passenger enplanement data by point of origin. U.S. Department of Transportation (DOT) Origin-Destination Passenger Survey is a comprehensive database that provides detailed trip and flight itinerary information for passenger travel on U.S. airlines. The data represent a 10% sample of all tickets that involve domestic travel on scheduled flights of U.S. airlines

These data were supplemented with data from the Department of Transportation T-100 database on the number of enplaned passengers for flights originating at a U.S. airport to an international destination.

Both passenger databases were downloaded from https://bit.ly/2P6N7SO (accessed March, 2019).

From these combined sources of data, we compiled state and temporal estimates by quarter of the total number of passengers enplaned. A total of 2.3 billion passengers were enplaned at their point of origin between 2012 to 2017.

### 2.2 Estimates of state prevalence of household firearm ownership, and state firearm-related laws

No official government statistics are available for household firearm ownership by state. However, firearm suicide rates relative to suicide rates by any means have been shown to be directly correlated to the prevalence of household firearm ownership [24–26]. We thus used suicide data from 2012 to 2017 by state and cause, obtained from the Centers for Disease Control (CDC) Wide-ranging ONline Data for Epidemiologic Research (WONDER) online database (http://wonder.cdc.gov/, accessed March, 2019), to estimate the state prevalence of household firearm ownership.

From Kalesan et al (2016) we determined the states that have child access prevention (CAP) laws (CA, CT, DE, FL, HI, IA, IL, KS, MA, MD, ME, MN, MT, NC, NH, NJ, NV, RI, TX, VA, and WI), and laws requiring safety training for purchase or licensing of firearms (CA, CT, HI, MA, MI, and RI) [27].

### 2.3 Statistical Methods

Over-dispersion is the presence of greater statistical variability in a data set compared to what would be expected based on a given statistical model, and is a common problem in statistical analyses of data in the life and social sciences [28]. In the following sub-sections, we describe the statistical modelling methods used in this analysis.

#### 2.3.1 Regression methods for over-dispersed count data

Count data are often over-dispersed relative to the Poisson model, in which case a more appropriate choice is the Negative Binomial model, which, like the Poisson distribution, includes the expected number of counts, *λ*, as a parameter, along with another parameter, *α* which is a measure of the amount of over-dispersion in the data [28]. The use of a Negative Binomial regression model when data are over-dispersed ensures that the estimates of the regression coefficients have accurate confidence intervals.

The Negative Binomial probability mass distribution for observing *k* counts when *λ* are expected is [28]

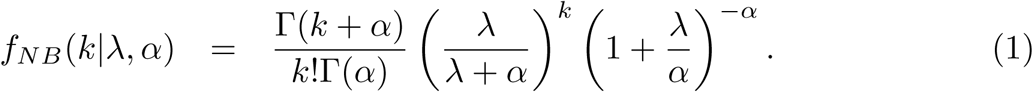

When *α* → ∞ the distribution approaches the Poisson distribution [28].

#### 2.3.2 Population standardisation of the Negative Binomial model

In a Poisson or Negative Binomial linear regression model we assume that

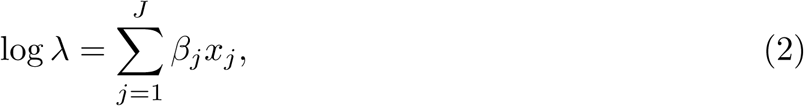

where *J* is the total number of regressors, *x*_*j*_. The best-fit parameters are the *β*_*j*_ which maximise the likelihood of observations. However, when analyzing the number of firearms detected per passenger, the number of firearms we detect in a particular locale will be proportional to the number of passengers in that locale, *N*. To take this into account, we perform what is known as population standardisation of the model [29], with

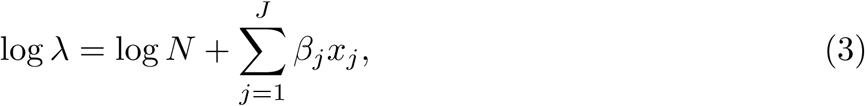

and perform the linear regression analysis with the coefficient of the log *N* term fixed to one.

We used population standardised Negative Binomial methods to linearly regress the logarithm of the national number of firearms detected per passenger on time, by quarter.

In a geospatial analysis, we also used population standardised Negative Binomial methods to linearly regress the logarithm of the number of firearms detected per passenger by state on the logarithm of the state firearm ownership prevalence, with additional factor levels for states with child access prevention laws, and states with laws requiring safety training for firearm purchase or licensing.

#### 2.3.3 Logistical regression methods for over-dispersed data

The Beta Binomial distribution is used to model over-dispersion in data that represent *k* successes out of *M* trials (for example *k* might be the number of firearms found loaded out of the total number, *M*, detected at TSA checkpoints over time). A key characteristic of the Beta Binomial distribution is that the variance is larger than that of the Binomial distribution with the same mean. It is important to take over-dispersion into account in order to obtain the correct confidence intervals on parameter estimates.

The Beta Binomial probability mass function, as a function of *k, M*, and two parameters *α* and *β* is [30]

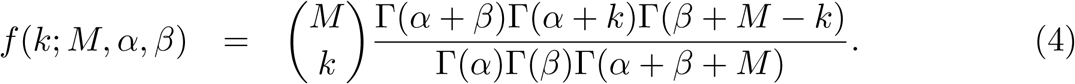

For the purposes of parameter estimation, it is convenient to express the PMF in terms of

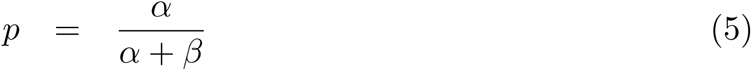

and

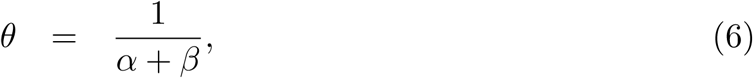

where *p* is the mean probability of success, and *θ* is a measure of the over-dispersion. When *θ* = 0 the distribution is the same as the Binomial distribution [30].

Here, in a temporal analysis, we used Beta Binomial logistic regression to linearly regress on time the logarithm of the odds that a firearm detected by the TSA was also found to be loaded. We also used Beta Binomial logistic regression to linearly regress on time the logarithm of the odds that a loaded firearm detected by the TSA was also found with a round chambered.

In addition, in a geospatial analysis, we used Beta Binomial logistic regression to model the logarithm of the odds, by state, that a firearm detected by the TSA was found to be loaded. Potential explanatory factor variables we examined were whether or not the state had child access prevention laws, and whether or not the state required safety training for firearm licensing. We also used Beta Binomial logistic regression to model the logarithm of the odds, by state, that a loaded firearm detected by the TSA was also found to have a round chambered, with the same explanatory variables.

## 3 Results

### 3.1 Temporal trends

The temporal trends in the number of firearms per passenger detected by the TSA nationally from 2012 to 2017 are shown in Figure 1. There has been a significant average relative rise of 14% per year in the firearms detected per passenger (population standardised Negative Binomial regression *p*< 0.001).

**Fig 1.**
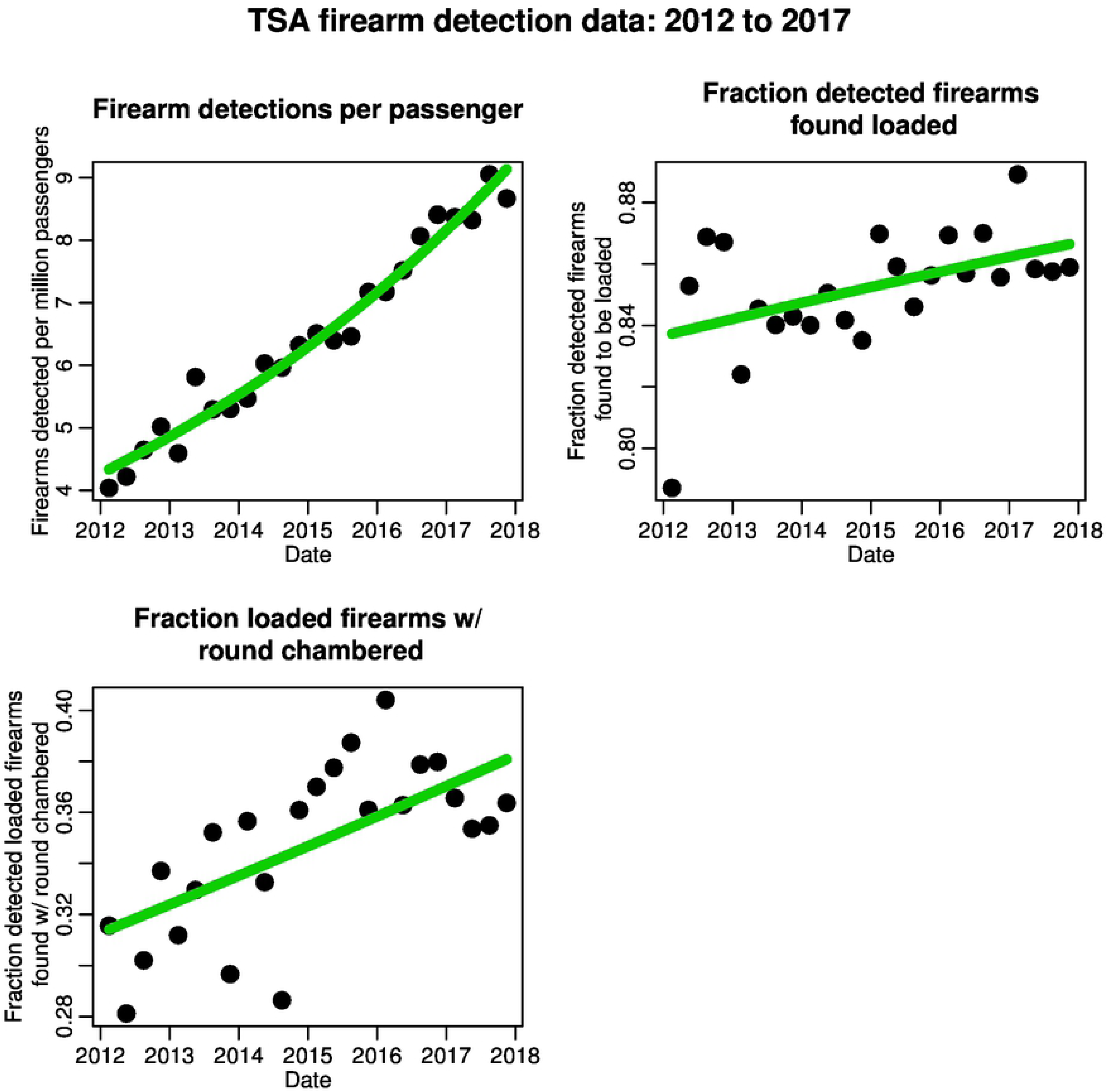
The top left plot shows the number of TSA firearm detections per million passengers at airports across the U.S. by quarter between 2012 to 2017. Overlaid in green are the results of the population standardised Negative Binomial linear regression in time. The top right plot shows the fraction of detected firearms that are found to be loaded by quarter, while the bottom left plot shows the fraction of detected loaded firearms that are also found to have a round chambered. Overlaid in green are the results of the Beta Binomial logistic linear regression in time.

Also shown in Figure 1 are the temporal trends in the fraction of firearms detected by the TSA that are found to be loaded. There has been a significant average relative annual rise of 4% in the odds of firearms being found loaded (Beta Binomial logistic regression *p* = 0.011).

The temporal trends in the fraction of loaded firearms that are also found to have a round chambered are also shown in Figure 1. There has been a significant average relative annual rise of 5% in the odds of loaded firearms to also have a round chambered (Beta Binomial logistic regression *p*< 0.001).

### 3.2 Geographical trends

In Figure 2, we show the average rate of firearm detections by passenger, the fraction of firearms found to be loaded, and the fraction of loaded firearms found to also have a round chambered, by geographical region in America. The highest rate of firearm detections per passenger is in the South, which also has the highest fraction of detected firearms that are found to be loaded (over 85%). The highest fraction of loaded firearms found to have a round chambered is in the Midwest (over 35%).

**Fig 2.**
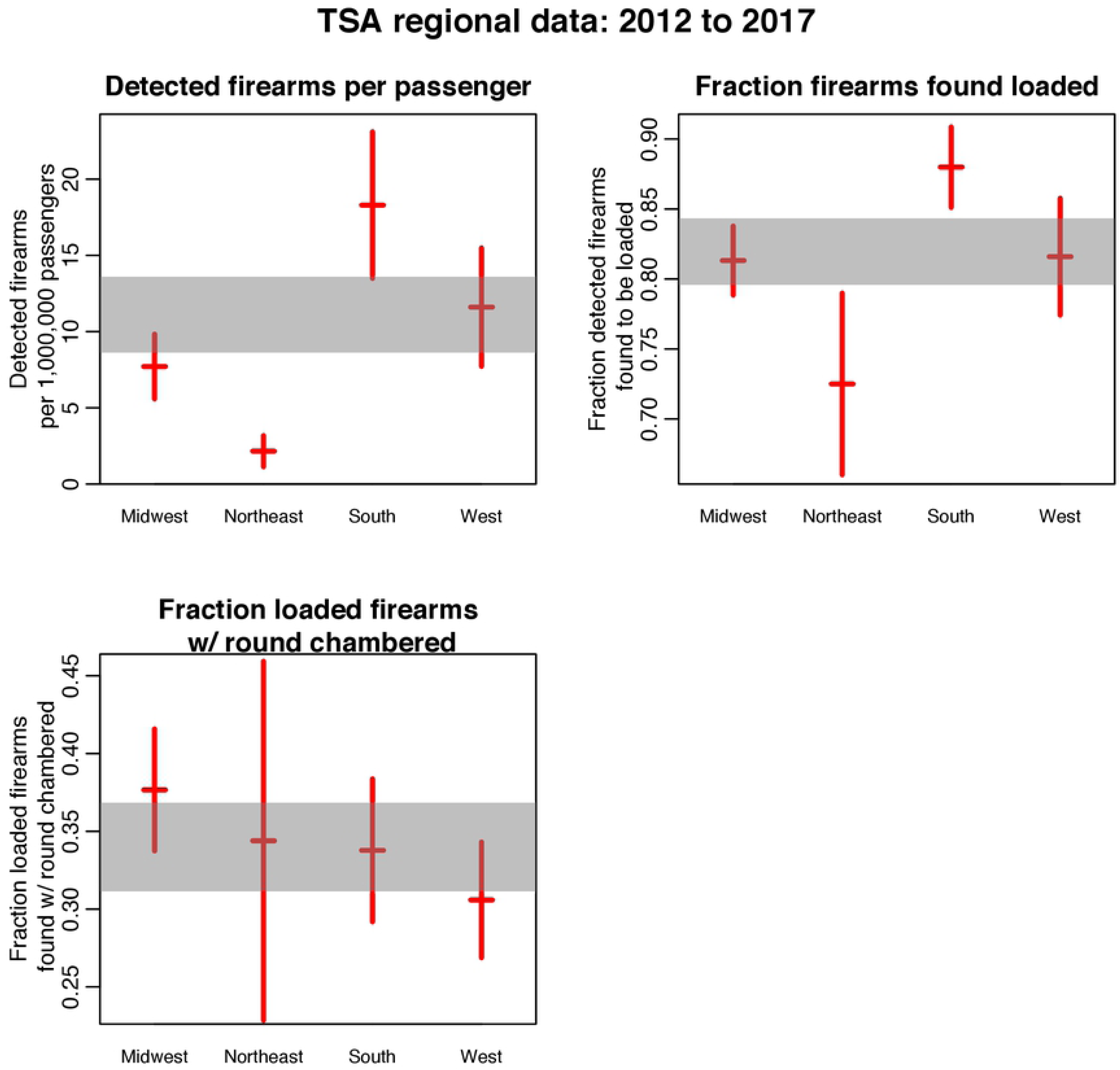
The average rate of firearm detections by passenger, the fraction of firearms found to be loaded, and the fraction of loaded firearms found to also have a round chambered, by the four geographical regions in America. The vertical bars represent the 95% confidence intervals, and the shaded gray area represents the 95% confidence interval for all data.

In Table 1 we show the results of population standardised Negative Binomial likelihood fits to the logarithm of the number of firearms detected per passenger, by state, regressing on the logarithm of state prevalence of household firearm ownership, and factors taking into account states that have child access prevention laws, and states that require safety training for purchase or licensing of firearms. As seen in Table 1, the rate of firearm detections by state has a significant positive relationship to prevalence of household firearm ownership, and a significant negative relationship to child access prevention laws and laws requiring safety training to purchase or license a firearm (*p <* 0.05 in all cases).

**Table 1.** Results of population standardised Negative Binomial likelihood fits to the logarithm of the number of firearms detected by the TSA per passenger, by state, regressing on the logarithm of state prevalence of household firearm ownership, and factors taking into account states that have child access prevention laws, and states that require safety training for purchase or licensing of firearms.

|  | Log prevalence<br>firearm ownership | Factor<br>CAP laws | Factor<br>safety training laws |
| --- | --- | --- | --- |
| Fit w/ only firearm ownership prevalence | $1.61 \pm 0.2$ ($p < 0.001$) | - | - |
| Fit w/ only CAP laws | - | $-0.95 \pm 0.23$ ($p < 0.001$) | - |
| Fit w/ only safety training laws | - | - | $-1.83 \pm 0.34$ ($p < 0.001$) |
| Fit w/ all effects | $1.16 \pm 0.2$ ($p < 0.001$) | $-0.5 \pm 0.18$ ($p = 0.005$) | $-0.82 \pm 0.28$ ($p = 0.003$) |

In Table 2 we show the results of Beta Binomial logistic fits to the logarithm of the odds, by state, of firearms detected by the TSA to also be found loaded, regressing on factors taking into account states that have child access prevention laws, and states that require safety training for purchase or licensing of firearms. As seen in Table 2, the dependent variable has a significant negative relationship to both factors (*p <* 0.05 in both cases).

**Table 2.** Results of Beta Binomial likelihood fits to the logarithm of the odds, by state, that firearms detected by the TSA were also found to be loaded, regressing on factors taking into account states that have child access prevention laws, and states that require safety training for purchase or licensing of firearms.

|  | Factor<br>CAP laws | Factor<br>safety training laws |  |
| --- | --- | --- | --- |
| Fit w/ only CAP laws | - | $-0.29 \pm 0.12$ ($p = 0.014$) | - |
| Fit w/ only safety training laws | - | - | $-0.47 \pm 0.17$ ($p = 0.010$) |
| Fit w/ both factors | $-0.24 \pm 0.11$ ($p = 0.039$) | $-0.39 \pm 0.17$ ($p = 0.032$) | |

In Table 3 we show the results of Beta Binomial logistic fits to the logarithm of the odds, by state, of loaded firearms detected by the TSA to also be found with a round chambered, regressing on factors taking into account states that have child access prevention laws, and states that require safety training for purchase or licensing of firearms. As seen in Table 3, the relationship to both factors was not significant (*p >* 0.05 in both cases).

**Table 3.** Results of Beta Binomial likelihood fits to the logarithm of the odds, by state, that loaded firearms detected by the TSA were also found to have a round chambered, regressing on factors taking into account states that have child access prevention laws, and states that require safety training for purchase or licensing of firearms.

|  | Factor<br>CAP laws | Factor<br>safety training laws |  |
| --- | --- | --- | --- |
| Fit w/ only CAP laws | - | $0.06 \pm 0.09$ ($p = 0.47$) | - |
| Fit w/ only safety training laws | - | - | $0.2 \pm 0.14$ ($p = 0.16$) |
| Fit w/ both factors | $0.04 \pm 0.09$ ($p = 0.65$) | $0.19 \pm 0.15$ ($p = 0.21$) | |

Beta Binomial logistic regression analysis with the state as a factor level to compare the fraction of detected firearms also found to be loaded to the national average revealed that the fraction was significantly higher than the national average in Alabama (136% higher than the national average, *p* = 0.020). The fraction was significantly lower than the national average in California, Hawaii, Illinois, Maryland, North Dakota, New Jersey, New York, and Rhode Island (*p <* 0.05 in all cases).

Beta Binomial logistic regression analysis with the state as a factor level to compare the fraction of loaded firearms also found to have a round chambered to the national average revealed that the fraction was significantly higher than the national average in California (31% higher than the national average, *p* = 0.044) and Indiana (54% higher, *p* = 0.019). The fraction was significantly lower than the national average in Montana (48% lower than the national average, *p* = 0.015) and Utah (62% lower, *p <* 0.001).

## 4 Discussion

In this analysis we examined firearms detected at TSA airport security checkpoints to estimate temporal and geospatial patterns in the storage and carry of firearms in unsafe arming configurations. While TSA detection efficiencies might have changed over time, the fraction of firearms found loaded, and the fraction found loaded that are also found to have a round chambered, are independent of these detection efficiencies and thus provide a useful means to assess temporal and geospatial trends in firearm storage and carry in these arming configurations, at least among the cross-section of firearm owners who are prone to negligent storage of firearms in unlocked luggage, or who are ignorant of federal laws that prohibit firearms in secure area of airports.

While representative of only a cross-section of firearm owners, these data provide a potential means to supplement the results of survey studies, and may help inform studies of the factors that underlie regional variations in firearm injury and mortality; survey studies can be expensive to undertake at the national level, and may also suffer from recall and social desirability bias [23], In addition, to the authors’ knowledge, ours is the first study to examine temporal and geospatial trends in the relative popularity of the practice of storing and carrying firearms with a round chambered, which is a particularly risky practice.

In the following subsections, we discuss the results of our temporal and geospatial analyses of the per passenger rate of TSA firearm detections, the fraction found loaded, and the fraction of loaded firearms additionally found with a round chambered.

### 4.1 Rate of TSA firearm detections per airline passenger

We found that the number of firearms detected per passenger has risen significantly at the national level since 2012. This can reflect either changing TSA screening practices, and/or changing firearm ownership prevalence, and/or changing storage habits, and/or increasing firearm carry. Further study will be needed to elucidate this further.

We found significant state variation in the rate of loaded firearms detected per passenger. There is a a significant relationship of this rate to the prevalence of firearm ownership. States with child access prevention laws also have significantly lower detection rates per passenger, as do states that require safety training for firearm purchase or licensing. This indicates that such laws appear to likely have some efficacy in discouraging unsafe storage practices (i.e. such as firearms stored unlocked in luggage), in-line with the conclusion of other studies based on survey data [31].

### 4.2 Fraction of firearms found to loaded

We found that the fraction of detected firearms found to be loaded has risen significantly nationally since 2012. This fraction is independent of changes in TSA screening efficiency because the efficiency affects both the numerator and denominator equally.

The detected firearms were a mixture of firearms negligently stored in luggage and forgotten, and firearms deliberately carried in to the airport, either in ignorance of federal law or in direct defiance of it. Stored firearms may or may not be loaded, but the firearms deliberately carried into the airport would likely have been. Because we do not know which firearms were carried into the airport deliberately, and which were simply stored and forgotten, it is difficult to assess whether the rise in the fraction of firearms found to be loaded reflects an increase in Americans storing firearms loaded, and/or whether it reflects an increase in Americans carrying firearms.

If the firearms detected by the TSA are dominated by negligently stored firearms, the upward trend we see in the fraction found to be loaded is in agreement with the results of a recent study that found that the fraction of children living in a home where a firearm is stored loaded and unlocked has risen more than two-fold since 2002 [23], despite the fact that the prevalence of household firearm ownership has been relatively static over that period [32].

The rising temporal trend in detection of loaded firearms that we observed in this study may be at least partly related to a shift in reasons for firearm ownership; a recent study has noted that the fraction of firearm owners stating that their primary reason for firearm ownership is self protection has risen from 26% to 48% between 1999 and 2013 [33], despite the fact that crime has decreased significantly over that period [34]. A previous study has shown that firearm owners who primarily own a gun for self defense are more likely to store their firearms loaded [35]. In addition, a recent non-peer-reviewed study has claimed that applications for concealed carry permits have risen dramatically in recent years [36].

The rise in owning guns for reasons of self defense may in part be due to public worries in the wake of recent high profile firearm violence in America, which has been implicated in a significant increase in handgun sales [37, 38], despite the fact the risk of being killed in such an event is rare [34]. Further study is needed to determine if local crime or high profile firearm violence is related to temporal trends in unsafe firearm storage practices, and/or firearm carry.

There is significant state variation of the fraction of firearms found to be loaded. We found the fraction is significantly lower in states with child access prevention laws, indicating that such laws appear to discourage the practice of storing firearms loaded, even when unlocked. In addition, we found that the fraction is significantly lower in states that require safety training for firearm purchase or licensing. When we compared the fraction of detected firearms found loaded by state to the national average, we found the fraction was significantly lower than the national average in California, Hawaii, Illinois, Maryland, North Dakota, New Jersey, New York, and Rhode Island (*p <* 0.05 in all cases); six out of these eight states have child access prevention laws.

We found that the fraction of detected firearms found loaded is highest in the South and lowest in the Northeast. This may again may reflect regional differences in unsafe storage practices (as noted in Section 2.2, states in the North are more likely than states in the South to have child access prevention laws and safety training laws for firearm licensing or purchase), and/or regional differences in the prevalence of firearm carry [39].

### 4.3 Fraction of loaded firearms found to also have a round chambered

We did not find systematic state variation in the fraction of loaded firearms that are also found to have a round chambered that is explainable by firearm storage laws or safety training requirements. However, this fraction has risen significantly nationally since 2012, again presenting a worrisome trend in storing or carry of firearms in very unsafe arming configurations.

When a firearm has a round chambered, if the safety is off (or the firearm does not have a safety), the trigger merely needs to be pulled to fire the weapon. For pistols, if a round is not chambered, one extra step must be taken to rack the slide to chamber a round before being able to fire the weapon.

The argument for storing or carrying a firearm with a round chambered is that removing the need to take the time to chamber a round is one less step that must be taken when using the firearm defensively. Like the rising trend we observe in the fraction of detected firearms that are found to be loaded, these patterns may be due to the increasing perception among the general public that firearms are primarily needed for self defense [33].

The practice of carrying or storing a firearm with a round chambered carries significant risk that is only mitigated by extensive training. For example, because they constantly face the risk of imminent threat, law enforcement officers generally carry with a round chambered [40], but the practice is considered so unsafe that the military generally only allows it when service members are in combat, or imminent risk of combat [41]. The practice of carrying a firearm with a round chambered in law enforcement and the military is a heavily regimented process due to the heightened safety concerns associated with such carrying configurations.

As with domestic law enforcement agencies, the military ensures that their members who are designated to carry loaded and chambered weapons are extensively trained to handle their weapons in this type of carrying configuration on a regular and documented basis [41]. Training has been shown to be effective in promoting safe storage practices [42], but a recent study has shown that almost 40% of civilian firearm owners have received no training on safe handling, safe storage, and injury prevention [43].

When we compared the state variation of the fraction of loaded firearms also found to have a round chambered to the national average, we found that one state in particular stood out; Utah had a fraction 62% lower than the national average (Beta Binomial logistic regression *p <* 0.001). Interestingly, Utah is the only state that mentions the issue of chambered rounds anywhere in its firearm laws; Utah allows open carry of a firearm without a permit as long as the firearm is at least two actions from being fired (e.g. for a revolver there must be no bullet in the chamber next to the firing pin, and for a pistol there user must have to rack the slide to chamber a round and then pull the trigger). Based on the results of our study, this Utah law has an apparent significant suppressive impact on the carry and storage of firearms with a round chambered.

Ours is the first study to examine the practice of carrying or storing firearms with a round chambered, and to raise the issue of the heightened risks this may pose to public health. Further study is needed to examine the reasons for the apparent significant rise in the practice of storing or carrying firearms with a round chambered, and to examine the effect this practice may have on firearm injury and mortality.

## 5 Summary

Our study used firearms detected at TSA airport security checkpoints between 2012 to 2017 to estimate temporal and geospatial trends in the storage and carry of firearms in unsafe arming configurations.

We found that the fractions of firearms found loaded, and loaded with a round chambered, have significantly increased since 2012. Both of these fractions are independent of TSA firearm detection efficiencies. The temporal and geospatial patterns we observe in the fraction of firearms found loaded are in broad agreement with prior studies based on survey data, and indicate that these data may be a useful adjunct in studies of patterns in firearm injury and mortality.

It should be cautioned that the firearms detected by the TSA represent a cross-section of firearm owners, and thus the findings of this study are not necessarily representative of trends in the broader firearm-owning population. However, the firearms detected, particularly the ones found with a round chambered, represent unsafe storage and carry practices, and thus even if these data only represent firearm owners found taking firearms through airport security, the upward temporal trends we observe in the prevalence of risky arming configurations is worrisome.

We found that state firearm child access prevention laws mitigate the fraction of firearms found loaded, but do not appear to mitigate the fraction of loaded firearms found with a round chambered. However, the Utah law allowing permitless open carry as long as the firearm does not have a round chambered has a significant suppressive effect on that unsafe arming configuration.

As of 2017, over 30% of firearms detected at airport security checkpoints by the TSA nationally are found with a round chambered. The safety issues posed by carrying or storing firearms with a round chambered have been overlooked in the academic literature, and our study underlines the need for further future research into the social determinants of this practice, and the risks the practice may pose for accidental injury.

## References

1. Fowler KA, Dahlberg LL, Haileyesus T, Annest JL. Firearm injuries in the United States. Preventive medicine. 2015;79:5–14.

2. Richardson EG, Hemenway D. Homicide, suicide, and unintentional firearm fatality: comparing the United States with other high-income countries, 2003. Journal of Trauma and Acute Care Surgery. 2011;70(1):238–243.

3. Grinshteyn E, Hemenway D. Violent death rates: the US compared with other high-income OECD countries, 2010. The American journal of medicine. 2016;129(3):266–273.

4. Fowler KA, Dahlberg LL, Haileyesus T, Gutierrez C, Bacon S. Childhood firearm injuries in the United States. Pediatrics. 2017; p. e20163486.

5. Bachier-Rodriguez M, Freeman J, Feliz A. Firearm injuries in a pediatric population: African-American adolescents continue to carry the heavy burden. The American Journal of Surgery. 2017;213(4):785–789.

6. Shah S, Hoffman RE, Wake L, Marine WM. Adolescent suicide and household access to firearms in Colorado: results of a case-control study. Journal of adolescent health. 2000;26(3):157–163.

7. Webster DW, Starnes M. Reexamining the association between child access prevention gun laws and unintentional shooting deaths of children. Pediatrics. 2000;106(6):1466–1469.

8. Miller M, Lippmann SJ, Azrael D, Hemenway D. Household firearm ownership and rates of suicide across the 50 United States. Journal of Trauma and Acute Care Surgery. 2007;62(4):1029–1035.

9. Miller M, Hemenway D, Azrael D. State-level homicide victimization rates in the US in relation to survey measures of household firearm ownership, 2001–2003. Social science & medicine. 2007;64(3):656–664.

10. Andrés AR, Hempstead K. Gun control and suicide: The impact of state firearm regulations in the United States, 1995–2004. Health Policy. 2011;101(1):95–103.

11. Conwell Y, Duberstein PR, Connor K, Eberly S, Cox C, Caine ED. Access to firearms and risk for suicide in middle-aged and older adults. The American Journal of Geriatric Psychiatry. 2002;10(4):407–416.

12. Shenassa ED, Rogers ML, Spalding KL, Roberts MB. Safer storage of firearms at home and risk of suicide: a study of protective factors in a nationally representative sample. Journal of Epidemiology & Community Health. 2004;58(10):841–848.

13. Miller M, Azrael D, Hemenway D, Vriniotis M. Firearm storage practices and rates of unintentional firearm deaths in the United States. Accident Analysis & Prevention. 2005;37(4):661–667.

14. Morgan ER, Gomez A, Rowhani-Rahbar A. Firearm Ownership, Storage Practices, and Suicide Risk Factors in Washington State, 2013–2016. American journal of public health. 2018;108(7):882–888.

15. Grossman DC, Mueller BA, Riedy C, Dowd MD, Villaveces A, Prodzinski J, et al. Gun storage practices and risk of youth suicide and unintentional firearm injuries. Jama. 2005;293(6):707–714.

16. Dowd MD, Sege RD, Gardner HG, Quinlan KP, Ewald MB, Ebel BE, et al. Firearm-related injuries affecting the pediatric population. Pediatrics. 2012;130(5):e1416–e1423.

17. Madenci AL. United States childhood gun-violence–Disturbing trends. In: 2013 AAP National Conference and Exhibition. American Academy of Pediatrics; 2013.

18. Santaella-Tenorio J, Cerdá M, Villaveces A, Galea S. What do we know about the association between firearm legislation and firearm-related injuries? Epidemiologic reviews. 2016;38(1):140–157.

19. Mechcatie E. Unsafe Firearm Storage in Homes with Children. AJN The American Journal of Nursing. 2018;118(6):17.

20. Tseng J, Nunõ M, Lewis AV, Srour M, Margulies DR, Alban RF. Firearm legislation, gun violence, and mortality in children and young adults: a retrospective cohort study. International Journal of Surgery. 2018;.

21. Okoro CA, Nelson DE, Mercy JA, Balluz LS, Crosby AE, Mokdad AH. Prevalence of household firearms and firearm-storage practices in the 50 states and the District of Columbia: findings from the Behavioral Risk Factor Surveillance System, 2002. Pediatrics. 2005;116(3):e370–e376.

22. Johnson RM, Miller M, Vriniotis M, Azrael D, Hemenway D. Are household firearms stored less safely in homes with adolescents?: Analysis of a national random sample of parents. Archives of pediatrics & adolescent medicine. 2006;160(8):788–792.

23. Azrael D, Cohen J, Salhi C, Miller M. Firearm storage in gun-owning households with children: results of a 2015 national survey. Journal of urban health. 2018;95(3):295–304.

24. Miller M, Azrael D, Hemenway D. Household firearm ownership and suicide rates in the United States. Epidemiology. 2002;13(5):517–524.

25. Miller M, Azrael D, Hepburn L, Hemenway D, Lippmann SJ. The association between changes in household firearm ownership and rates of suicide in the United States, 1981–2002. Injury Prevention. 2006;12(3):178–182.

26. Azrael D, Cook PJ, Miller M. State and local prevalence of firearms ownership measurement, structure, and trends. Journal of Quantitative Criminology. 2004;20(1):43–62.

27. Kalesan B, Mobily ME, Keiser O, Fagan JA, Galea S. Firearm legislation and firearm mortality in the USA: a cross-sectional, state-level study. The Lancet. 2016;387(10030):1847–1855.

28. Lloyd-Smith JO. Maximum likelihood estimation of the negative binomial dispersion parameter for highly overdispersed data, with applications to infectious diseases. PloS one. 2007;2(2):e180.

29. Osgood DW. Poisson-based regression analysis of aggregate crime rates. Journal of quantitative criminology. 2000;16(1):21–43.

30. Hughes G, Madden L. Using the beta-binomial distribution to describe aggregated patterns of disease incidence. Phytopathology (USA). 1993;7.

31. Prickett KC, Martin-Storey A, Crosnoe R. State firearm laws, firearm ownership, and safety practices among families of preschool-aged children. American journal of public health. 2014;104(6):1080–1086.

32. Smith TW, Son J. Trends in gun ownership in the United States, 1972–2014. General Social Survey Final Report Chicago: University of Chicago: NORC. 2015;.

33. Center PR. Why own a gun? Protection is now top reason; 2013.

34. Bagalman E, Caldwell SW, Finklea KM, McCallion G. Public Mass shootings in the United States: Selected implications for federal public health and safety policy. Congressional Research Service. 2013;.

35. Weil DS, Hemenway D. Loaded guns in the home: analysis of a national random survey of gun owners. Jama. 1992;267(22):3033–3037.

36. Lott JR, Whitley JE, Riley R. Concealed carry permit holders across the United States. Social Science Research Network electronic library. 2015;2015(online):ID–2629704.

37. Wallace LN. Responding to violence with guns: Mass shootings and gun acquisition. The Social Science Journal. 2015;52(2):156–167.

38. Studdert DM, Zhang Y, Rodden JA, Hyndman RJ, Wintemute GJ. Handgun acquisitions in California after two mass shootings. Annals of internal medicine. 2017;166(10):698–706.

39. Felson RB, Pare PP. Gun cultures or honor cultures? Explaining regional and race differences in weapon carrying. Social forces. 2010;88(3):1357–1378.

40. Sullivan GG, Hamilton GM. Carrying of Firearms and Use of Force for Law Enforcement and Security Duties. 1993; Army regulation 190-14:1–10.

41. Pink CK, Ornelas MG. Arming and use of force by Air Force personnel. 2012;Air Force Instruction Manual 31-207:1–39.

42. Crifasi CK, Doucette ML, McGinty EE, Webster DW, Barry CL. Storage practices of US gun owners in 2016. American journal of public health. 2018;108(4):532–537.

43. Rowhani-Rahbar A, Lyons VH, Simonetti JA, Azrael D, Miller M. Formal firearm training among adults in the USA: results of a national survey. Injury prevention. 2017;doi:10.1136//injuryprev-2017-042352.

